# Motor axon excitability measures in the rat tail are the same awake or anaesthetized using sodium pentobarbital

**DOI:** 10.1101/651927

**Authors:** Kelvin E Jones, David J Bennett

## Abstract

**Background:** Nerve excitability tests in sciatic motor axons are sensitive to anaesthetic choice. Results using ketamine/xylazine (KX) are different from those using sodium pentobarbital (SP). It is not clear which results are most similar to the awake condition, though results using SP appear more similar to human results.

**Methods:** Nerve excitability in tail motor axons was tested in 8 adult female rats with a chronic sacral spinal cord injury. These animals have no behavioural response to electrical stimulation of the tail and were tested awake and then anaesthetized using SP.

**Results:** The nerve excitability test results in the awake condition were indistinguishable from the results when the same rats were anaesthetized with sodium pentobarbital. Summary plots of the test results overlap within the boundaries of the standard error and paired t-tests on the 42 discrete measures generated by nerve excitability testing yielded no significant differences (after Bonferroni correction for multiple comparisons).

**Conclusions:** Nerve excitability test results in rat motor axons are the same whether the animals are awake or anesthetized using sodium pentobarbital.

## Background

The electrophysiological properties of peripheral axons can be assessed using an *in vivo* rat model to measure axonal excitability using the technique of threshold tracking [1]. The same methods have been used clinically to assess the pathophysiology of peripheral nerves (e.g. [2, 3]). Recent studies from our lab have used this methodology to measure differences between motor axons innervating fast-compared to slow-twitch muscle [4, 5]. We frequently use an intraperitoneal injection of a cocktail comprised of ketamine and xylazine (KX) for acute procedures lasting about an hour and test withdrawal reflexes regularly, supplementing with additional injections of KX if the reflex becomes evident [6]. In Canada, ketamine is also used for purposes other than small animal surgery [7] and therefore if you run out, there can be a delay in filling out the appropriate paperwork to procure more ketamine to continue experiments.

This was the situation we found ourselves in and, rather than wait, we turned to an older alternative veterinary anesthetic, sodium pentobarbital [8]. After all, we were testing the peripheral nervous system and neither anesthetic: KX nor SP, had contraindications for the electrophysiological properties of peripheral nerves, or so we thought. Data collection was nearly complete for a Masters student project [4] and we thought perhaps we could collect data from a few animals substituting SP for KX as the anesthetic. Wrong … the initial results looked so different we decided not to continue data collection using SP and wait for our ketamine to arrive to complete the project. However the data from the sciatic nerve using KX appeared to look quite different compared to data from motor axons in the tail using KX, especially during the recovery cycle test [1].

Eventually we decided to redo the experiments using SP to determine if the initial results suggesting an anesthetic effect were spurious or if nerve excitability test results could be confounded by anesthetic choice. The results were not spurious and it was clear that the data gathered using SP *appeared* more similar to what we had come to expect, especially during the recovery cycle [5]. However, similarity to expectations is perhaps not the most rigorous method to use in science.

Ideally, when measuring axonal excitability in rodents an effect of anesthetic is to be avoided since the goal is to provide data similar to the awake human testing condition. The chronic sacral spinal injury rat model provided a unique opportunity to test animals in the awake condition since they can not sense a stimulus to their tail [9]. We decided to perform the nerve excitability test in these animals, awake and then under general anesthesia to measure anaesthetic effects in the same animal.

## Methods

The procedures were approved by the University of Alberta’s Animal Care and Use Committee (ACUC): Health Sciences #2 (AUP00000733) and were in accordance with the Canadian Council on Animal Care Guidelines. Our animal protocol allowed for use of one intraperitoneal anaesthetic and we chose SP to determine if there were differences from the awake condition.

### Rats

Thirteen female Sprague–Dawley rats, weighing an average of 417 g and average age of 50-weeks (standard deviation 4.8 weeks), were made available for these experiments. These animals were part of an ongoing research program in Dr Bennett’s lab examining the pathophysiology and treatment of spasticity following spinal cord injury [9, 10, 11]. All animals had a complete sacral spinal transection (at S2) at about 2 months of age and were tested in these experiments more than 300 days following the initial injury. The animals had come to the end of the follow-up period for Dr Bennett’s research and were scheduled to be euthanized.

### Nerve Excitability Test (NET)

Tail motor axons were tested using the NET as previously described [1, 4, 5] using the QtracW threshold tracking software https://digitimer.com/products/research-electrophysiology/software-updates/qtracw-threshold-tracking-software/. The rats were initially tested in the awake condition by placing them in a plexiglass tube to restrain them while leaving their tail exposed for testing (Figure 1). Data collection took about 12 minutes, after application of electrodes for stimulation and recording the compound muscle action potential (CMAP) from the tail. Afterwards the animal was: removed from the restraining tube, given an intraperitoneal injection of sodium pentobarbital (40 mg/kg) and retested when it reached a surgical plane of anaesthesia (determined by absence of withdrawal and corneal reflexes, similar to [5]). In the anesthetized condition the temperature of the tail was measured with a thermocouple and the rat - including tail - were placed on a Deltaphase isothermal warming pad https://www.braintreesci.com/products.asp?dept=128 (Figure 2). Temperature was maintained during anaesthesia at the same temperature recorded in the awake condition, an average of 33.5 degC (SD ± 1.2). Following data collection the rats were euthanized by anaesthetic overdose and cervical dislocation. Data collection was **not** successful in all 13 animals: 1 animal was too large for the restraint tube, 3 animals could not be tested because the stimuli during the NET generated long-duration muscle spasms, and technical issues prevented repeat data collection from one animal. Repeated measures of the NET were successful in 8 animals.

**Figure 1.**
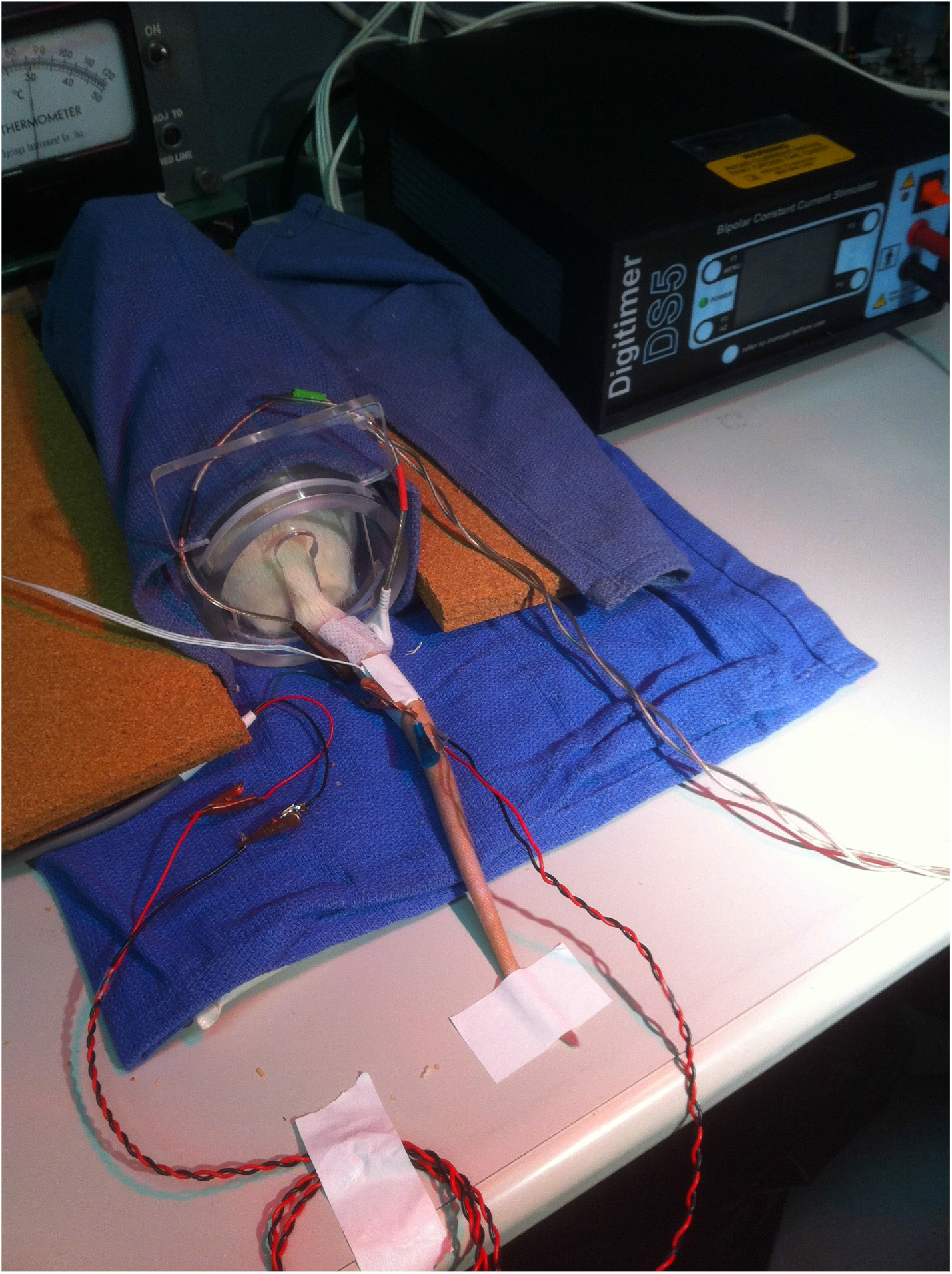
NET in Awake Condition. A rat in the restraining tube during testing. Draping the tube with towels and working in a dim environment was essential to keep the rats calm and quiet during the NET.

**Figure 2.**
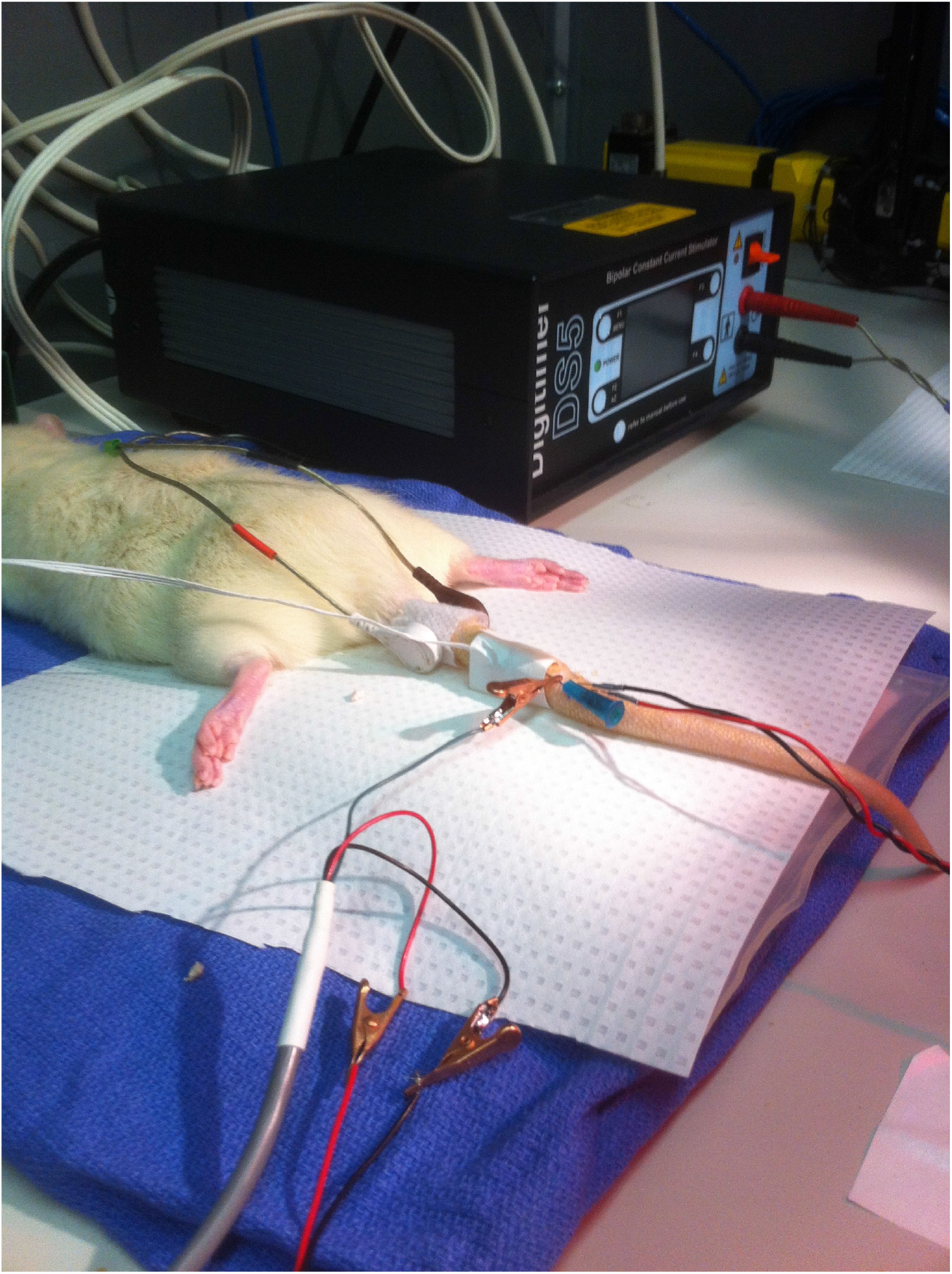
NET in Anesthetized Condition. The stimulating electrodes were in the same position and the subdermal electrodes for recording CMAP were placed in the same area as during the previous test in the awake condition.

### Analysis

Data analysis and visualization were done using QtracW software. The time-series waveform data are presented as means (solid) and standard error of the mean (dashed) for the Awake (red) and SP (blue) conditions. (**Note:** Plotting the standard error, rather than standard deviation, was done to visually detect any potential regions where estimates of the means did not overlap, thus facilitating the possibility of seeing differences in the two conditions.) Paired t-tests on the 42 dependent variables generated by the NET were done and a Bonferroni correction applied to correct for multiple comparisons: desired *α* = .05 is ≡ .00119. (Though correction was not needed as all p-values ranged from a low of .052 (TEh20 10-20 ms) to .941 (TEh, peak −70%) with an average of .469 and median value of .329, *i.e*. all values are greater than the somewhat arbitrary threshold of .05 even without correction.) The figure legends report mean (± standard deviation) for some associated discrete measures.

## Results

The data from the same rats, awake and anesthetized with SP were indistinguishable. This was true for all the components of a full NET: Charge-Duration (QT, Figure 3), Threshold Electrotonus (TE, Figure 4), Current-Threshold (IV, Figure 5), and Recovery Cycle (RC, Figure 6).

**Figure 3.**
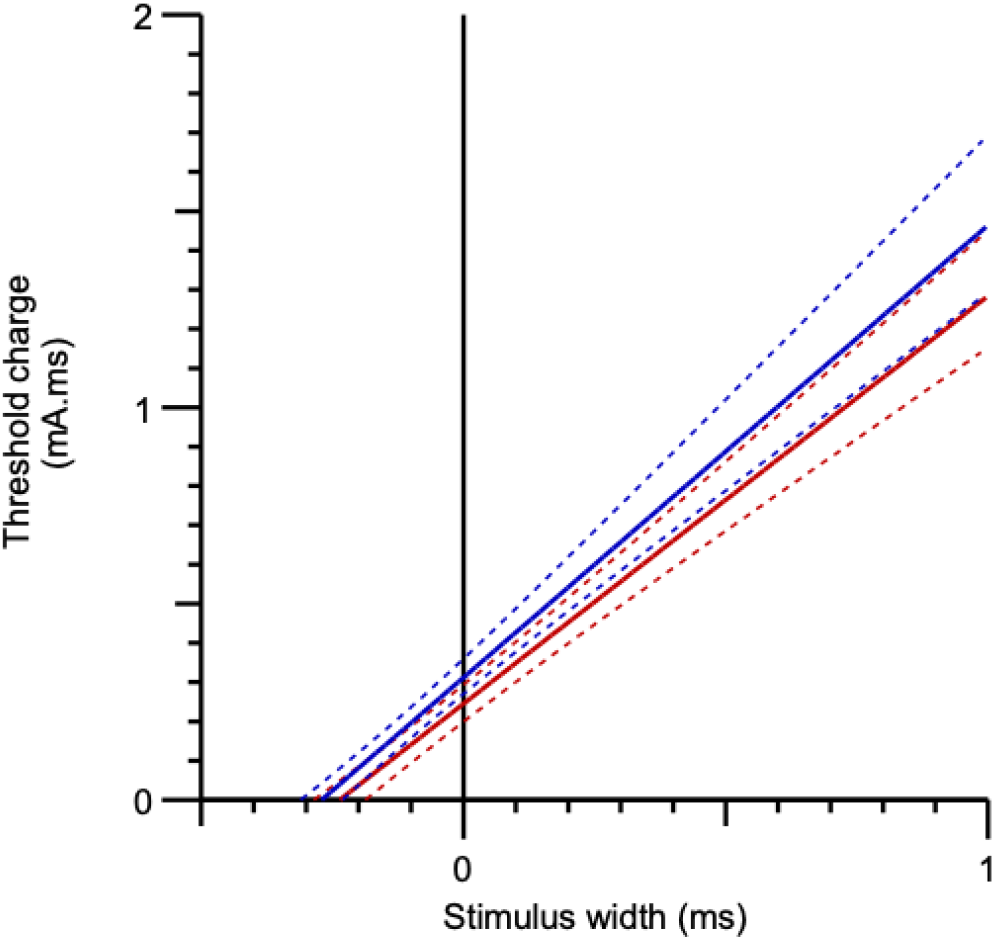
Charge-Duration. The rheobase was 0.996 ±1.4 for the Awake condition (red) and 1.081 ±1.5 in the SP condition. Strength-duration time constants were 0.252 ±0.139 and 0.289 ±0.120 in the Awake and SP conditions respectively.

**Figure 4.**
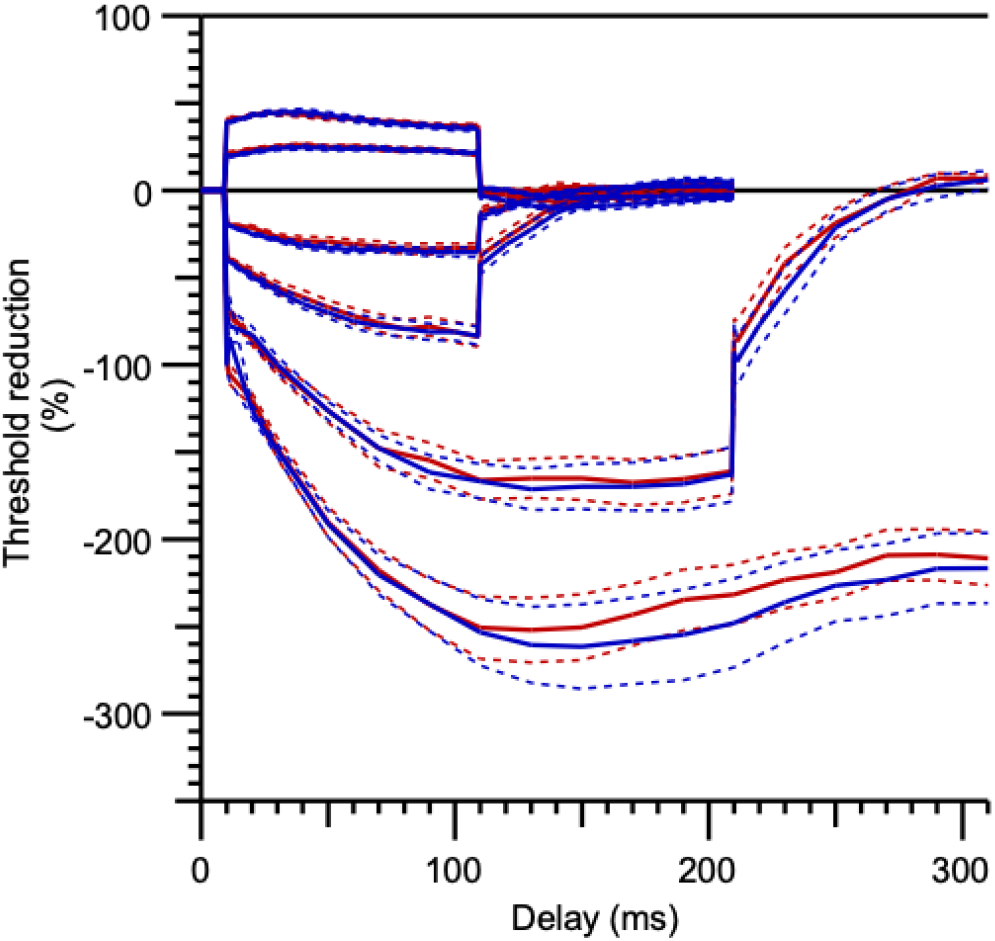
Threshold Electrotonus. Mean values in the Awake (red) and SP (blue) conditions completely overlapped during the standard 20% and 40% depolarizing and hyperpolarizing conditions. The peak threshold reductions during the 300 ms, 100% hyperpolarizing conditioning had a mean difference of 54.07 ±38.4, with smaller size in the Awake condition.

**Figure 5.**
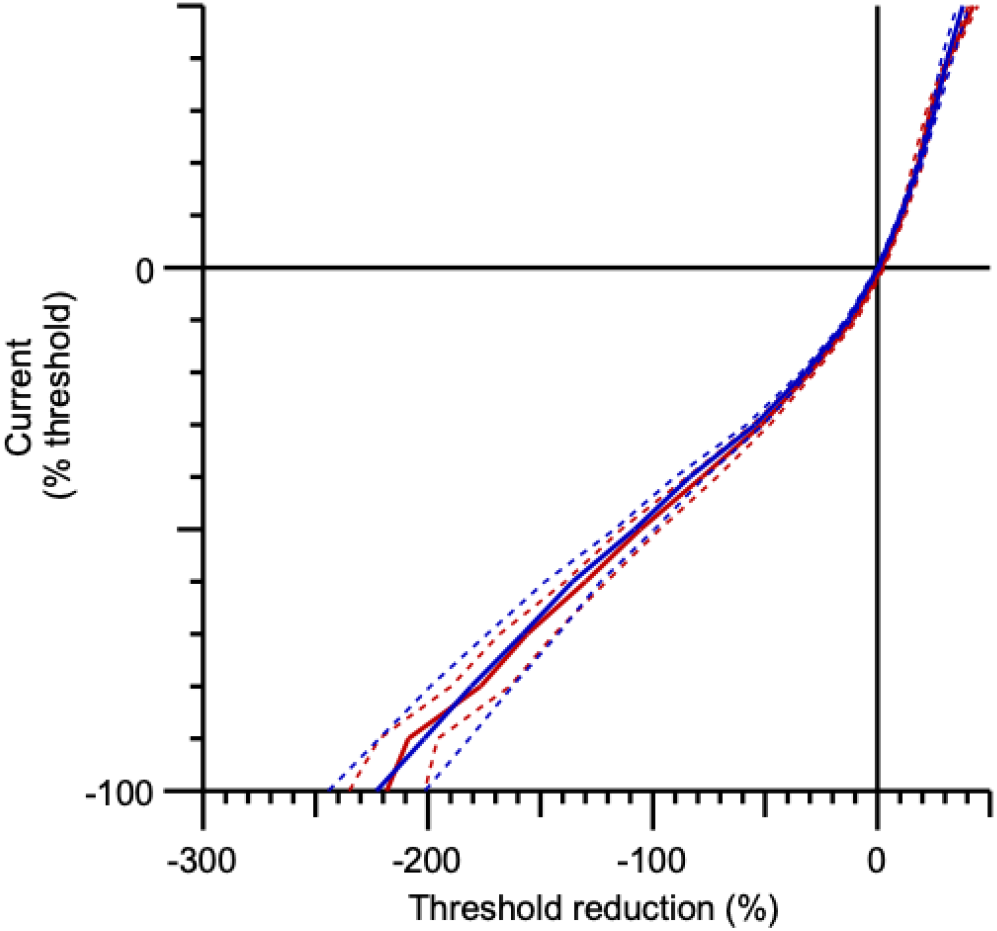
Current-Threshold. Mean values in the Awake (red) and SP (blue) conditions overlapped over the full range of 200 ms depolarizing and hyperpolarizing conditioning currents. There were no differences in the resting, minimum or hyperpolarized I/V slopes: (mean differences) .0107, .0337, and -.0025 respectively.

**Figure 6.**
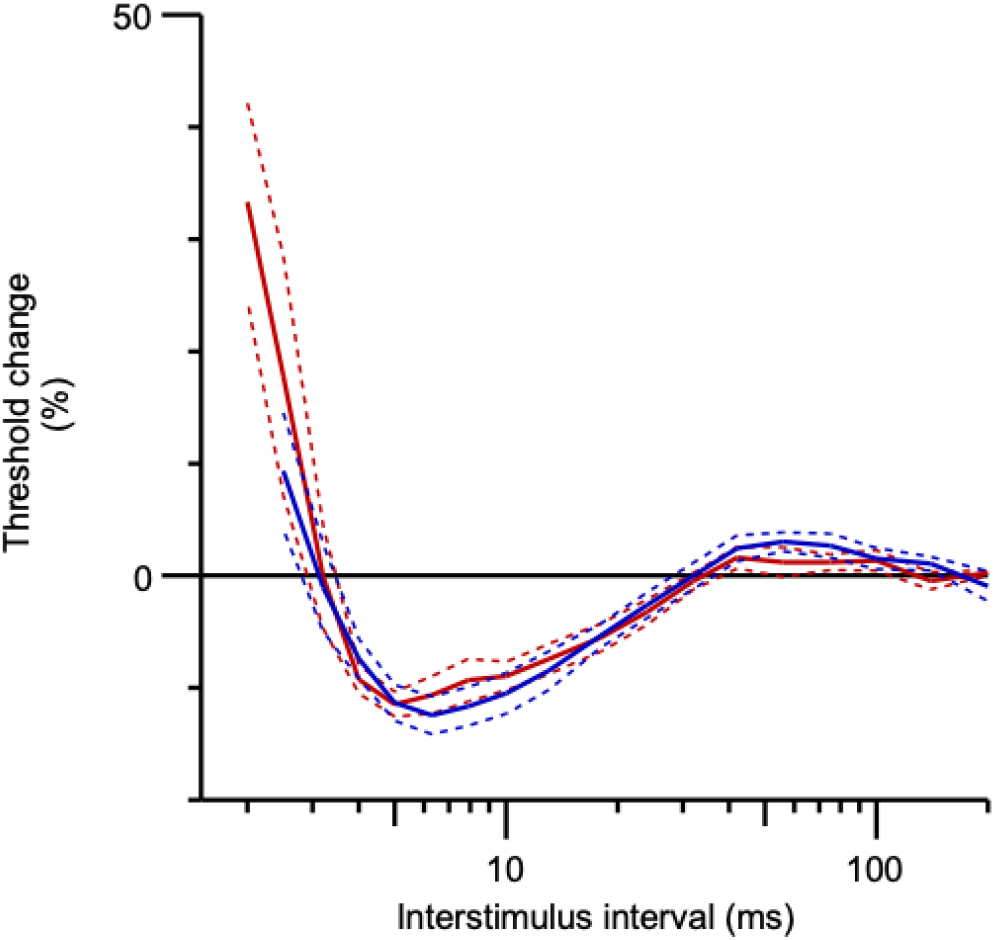
Recovery Cycle. The relative refractory period in the Awake (red) and SP (blue) conditions were similar: 2.92 ±1.15, 3.13 ±1.19. Threshold change during the superexcitable periods were similar: −10.98 ±3.29 (Awake) & −12.08 ±4.13 (SP). During the late subexcitable period average thresholds in the two conditions were: 2.31 ±1.45 & 3.53 ±2.11.

## Discussion

It is clear that there is no overt effect of SP anaesthesia on nerve excitability measures from rat tail motor axons compared to the awake condition. We believe this finding can be generalized to testing of motor axons in the sciatic nerve, though that nerve can not be tested in the awake condition since the surgical procedure must be done under general anaesthesia. The difference in results using KX versus SP, together with these results comparing Awake versus SP in the same animal, support our assertion that when comparing fast-verus slow-axons use of SP is preferred to KX [5].

### Limitations

- The female animals used here are much older than those used in the studies examining fast and slow axons in the sciatic nerve. The current animals are on average 50 weeks of age (*i.e*. nearly 12 months, or about 350 days old) compared to young, but sexually mature females of about 13 weeks of age (*i.e*. about 3 months, or at least 90 days old) [4, 5]. Since there are maturation effects that are evident when testing rodent axon excitability [12, 13], it would have been ideal to have tested animals of similar ages. However, we are not comparing excitability measures across ages, but rather the effects of anesthetic choice, so the difference in ages is less relevant.
- All the animals in this study had a chronic spinal cord injury (SCI). Again, we are not comparing absolute excitability values in the tail motor axons of old female rats with a SCI to sciatic motor axons of younger female rats without a SCI. It remains to be demonstrated how nerve excitability from tail motor axons compares to fast- and slow-motor axons in the leg in the same animals. The potential problem is whether changes in the motor axons of the tail following a chronic SCI lead to an insensitivity to SP anesthetic; meaning the lack of effect seen here is a result of the SCI and would be different in intact animals. While possible, we believe this is an unlikely scenario.

### Implications and Questions

The opportunity to measure axon excitability in animals with a chronic spinal cord injury generated many additional questions beyond the immediate goal of measuring an anesthetic effect compared to the awake condition. First, we chose to use SP in these experiments to be consistent with the approved animal protocols. It may be interesting to confirm whether tail motor axons are susceptible to the use of KX similar to the fast- and slow-axons of the leg. An attempt to examine this question was made by plotting data from young, female rats with intact spinal cords [1] and the present data (Figure 7).

**Figure 7.**
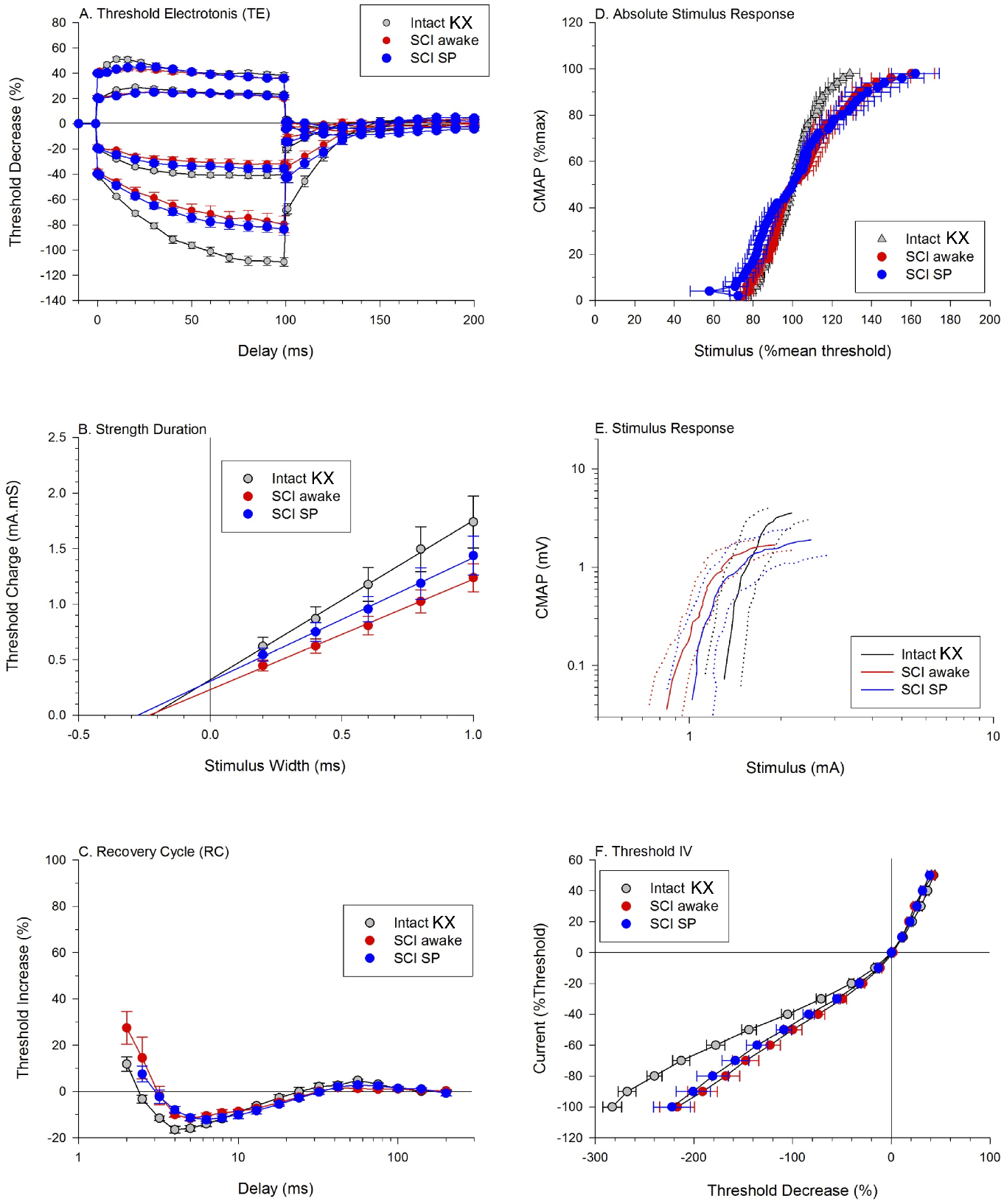
Comparing Tail Motor Axons: (Young, KX, Intact) versus (Older, SP, SCI). The data previously published in [1] were made available to examine alongside the current data. There are surprisingly fewer differences than expected given the differences in: age, anesthesia, and injury. There is an increased threshold in the Intact KX data illustrated in the increased rheobase (i.e. slope, panel B) and rightward shift of the stimulus response curve (panel E). There are greater changes in threshold with hyperpolarizing conditioning stimuli (panels A and F) that are consistent with reduced efficacy of H-current in the Intact KX data. The refractory period is shorter and superexcitability has a greater threshold change. Data plotted as means and standard errors.

It would also be of interest to compare tail motor axons of intact animals to animals with a chronic sacral transection injury to determine if axonal excitability measures change after a spinal cord injury. There are changes in these measures in people following a spinal cord injury [14] and those changes can be ameliorated with short-term nerve stimulation to induce activity in the motor axons [15]. In the Bennet lab, the chronic spinal rats can be divided into two categories: those who go on to develop tail-muscle spasticity (~ 95%) and those who do not and have nonspastic, inactive tails [11]. The data presented here are from the former group of rats that developed spasticity. We hypothesize that chronic spinal rats that have nonspastic, inactive tails would show changes similar to people without spasticity while the spastic rats should show less changes, in accordance with the effects of exogenous stimulation in people following a SCI [14, 15].

## List of Abbreviations

KX: ketamine-xylazine
SP: sodium pentobarbital
NET: Nerve Excitability Test
SCI: spinal cord injury

## Declarations

### Ethics approval

The procedures were approved by the University of Alberta’s Animal Care and Use Committee (ACUC): Health Sciences #2 (AUP00000733) and were in accordance with the Canadian Council on Animal Care Guidelines.

### Consent for publication

Not applicable

## Acknowledgements

I am grateful to Neil Tyrem an for his expert technical assistance in the lab and for instigating the discussion with Leo Sanelli from Dr Bennett’s lab about the possibility of pursuing these experiments. Thanks to Professor Hugh Bostock for sharing the data from [1] in Figure 7.

